# Oxytocin differentially modulates amygdala responses during top-down and bottom-up aversive anticipation

**DOI:** 10.1101/2020.03.14.992172

**Authors:** Fei Xin, Xinqi Zhou, Debo Dong, Zhongbo Zhao, Xi Yang, Qianqian Wang, Yan Gu, Keith M. Kendrick, Antao Chen, Benjamin Becker

## Abstract

The ability to successfully regulate negative emotions such as fear and anxiety is vital for mental health. The neuropeptide oxytocin (OXT) acts as important modulator of emotion regulation, as reflected by reduced amygdala responses but increased amygdala–prefrontal cortex (PFC) functional connectivity in response to threatening stimuli. The present randomized, between-subject, placebo (PLC)-controlled pharmacological study combined intranasal administration of OXT with functional MRI during an explicit (cognitive) emotion regulation (i.e. distancing reappraisal) paradigm in 65 healthy male participants to investigate the modulatory effects of OXT on both bottom-up and top-down emotion regulation. OXT attenuated the activation in posterior insular cortex and amygdala during anticipation of top-down regulation of predictable threat stimuli in participants with high trait anxiety, providing evidence to support the anxiolytic action of OXT. In contrast, OXT enhanced amygdala activity during bottom-up anticipation of an unpredictable threat stimulus in participants with low trait anxiety. OXT may thus facilitate top-down goal-directed attention by attenuating amygdala activity in high anxiety individuals, while promote bottom-up attention/vigilance to unexpected threat by enhancing anticipatory amygdala activity in low anxiety individuals. The opposite effects of OXT on anticipatory amygdala activation in high versus low anxiety individuals may suggest a baseline anxiety level dependent mechanism via which OXT promotes optimal levels of amygdala activation during the anticipation of an imminent threat. OXT may thus have the potential to promote an adaptive balance between bottom-up and top-down attention systems depending on individual levels of pre-treatment trait anxiety levels.

## Introduction

From an evolutionary perspective, fear and anxiety elicit defensive responses to detect, react and cope with imminent or potential threat and to avoid harm (Steimer, 2002). Both, excessive as well as deficient anxiety and fear strongly interfere with the ability to react to and cope with potential threat and life challenges, as exhibited in anxiety disorders or brain lesion patients (Becker et al., 2012; Greenberg et al., 1999). The ability to successfully regulate negative emotions such as fear and anxiety is vital for mental health, well-being, and social functioning (Gross, 2014; Gross and John, 2003). Consequently dysregulations in the fine-grained interplay between the generation and regulation of negative emotions represent transdiagnostic deficits across major psychiatric disorders (Sloan et al., 2017; Zilverstand et al., 2017). Behavioral interventions (Campbell-Sills and Barlow, 2007; Gross et al., 2006; Jazaieri et al., 2015), neurofeedback targeting the amygdala-prefrontal circuits (Cohen Kadosh et al., 2016; Zhao et al., 2019) and pharmacological agents primarily targeting serotonergic and GABA-ergic neurotransmission (Farach et al., 2012; Singewald et al., 2015) can facilitate the regulation of negative emotions, including anxiety and fear. However, not all patients respond adequately to the currently available treatment strategies and the pharmacological interventions may induce negative side-effects which often limit compliance with the treatment protocols, thus novel strategies to improve emotion regulation are urgently needed.

Emotion generation and regulation involves the interaction of intrinsic/automatic bottom-up processes and controlled/deliberately top-down processes (Ochsner and Gross, 2005; Ochsner and Gross, 2007; Suri et al., 2013). Bottom-up processes are stimulus-driven and initiated by salient stimuli in the environment. In contrast, top-down processes are goal-driven and context-dependent (Beck and Kastner, 2009; Desimone and Duncan, 1995; Sussman et al., 2016a). During both the (pre-stimulus) anticipation and advent of the aversive events, the bottom-up and top-down processes rely on differential neural systems and temporal dynamics. Bottom-up emotional processes are mediated by subcortical systems such as the amygdala and hypothalamus. In contrast, top-down regulatory processes rely on prefrontal executive control systems (Comte et al., 2016; Ochsner et al., 2009). Relative to the automatic stimulus-driven bottom-up processes, top-down processes take into account the context of threat (predictive) cues and initiate anticipatory goal-directed responses to facilitate threat-vigilance and emotion regulation. Although anticipatory cues or goals can induce more anxiety, they may also contribute to perceptual and attentional prioritization of threatening stimuli such that anticipatory cues can accelerate stimulus coding in the visual and attention systems (Hahn and Gronlund, 2007; Mazzucato et al., 2019; Sussman et al., 2016a; Yoshida and Katz, 2011).

The anticipatory response to low imminent, uncertain or future threat specifically refers to anxiety whereas fear is elicited by a high imminent and certain threat (Davis et al., 2010; Grupe and Nitschke, 2013; Tovote et al., 2015). The neural circuits that mediate fear-associated responses and the implicit and explicit regulation of imminent threat (e.g. Etkin et al., 2015) as well as dysregulations in these processes in psychiatric disorders (e.g. Zilverstand et al., 2017) have been extensively studied. However, despite the important role of exaggerated threat anticipation and impaired regulation of the anticipatory response for the pathophysiological mechanism of anxiety disorders (Nitschke et al., 2009; Sussman et al., 2016b), only few studies focused on the pre-stimulus anticipatory period. Previous studies indicate that - In line with the proposed pathological mechanism in anxiety disorders - pre-stimulus anticipatory brain activity is predictive of subsequent emotion regulation success or failure (Denny et al., 2014).

Previous animal models and human studies provided extensive evidence for an important role of the hypothalamic neuropeptide oxytocin (OXT) in anxiety, and based on these findings intranasal OXT has been suggested as potential promising pharmacological treatment for anxiety and fear-related disorders (Eckstein et al., 2015; Knobloch et al., 2012; Neumann and Slattery, 2016; Spengler et al., 2017). Converging evidence indicates an anxiolytic potential of intranasal OXT reflected by reduced amygdala responses but increased amygdala–prefrontal cortex (PFC) functional connectivity in response to threatening stimuli in healthy adults (Domes et al., 2007; Eckstein et al., 2015; Petrovic et al., 2008; Striepens et al., 2012). Two of the key regions modulated by OXT, PFC and amygdala, play crucial roles in emotion generation and regulation (Banks et al., 2007; Etkin et al., 2015; Lee et al., 2012). In addition to effects on this circuitry in response to imminent threat stimuli, intranasal OXT has been repeatedly demonstrated to enhance intrinsic amygdala-prefrontal connectivity in the absence of threat-related stimuli, suggesting potential effects on the fear and anxiety related circuitry that precede the actual encounter of threat (Eckstein et al., 2017; Sripada et al., 2013). The effects of OXT on human cognition and behavior are partly mediated through widely distributed central OXT receptors, with particularly dense expression of OXT-sensitive receptors in the neural systems associated with emotion generation and regulation, including the prefrontal and anterior cingulate cortex, as well as the amygdala, hippocampus, striatum and the brainstem (Bethlehem et al., 2017; Boccia et al., 2013; Quintana et al., 2019). Based on these findings, we expected that OXT may modulate the responses of anticipation of threat in amygdala and regulation of anticipatory threat in the prefrontal cortex.

Against this background, the present randomized, between-subject, placebo (PLC)-controlled pharmacological fMRI study combined the intranasal administration of OXT with functional MRI during an explicit (cognitive) emotion regulation (i.e. distancing reappraisal) paradigm in 65 healthy male participants to investigate the modulatory effects of OXT on both bottom-up and top-down emotion regulation. To this end we employed an explicit emotion regulation paradigm (i.e. cognitive reappraisal) which allows to study bottom-up and top-down emotion generation and regulation simultaneously. The cognitive reappraisal paradigm includes i) a ‘Look’ condition during which participants are instructed to simply view a threatening image and react naturally (bottom-up trials), and ii) a ‘Distance’ condition during which participants are instructed to regulate their emotional response by means of cognitive reappraisal (top-down trials) (Goldin et al., 2008; Koenigsberg et al., 2009; Ochsner et al., 2004). Moreover, this paradigm additionally includes a pre-stimulus anticipatory period between a cue signaling the instruction for the subsequent threat stimulus (‘Look’, ‘Distance’), which allows to examine anticipatory responses to an uncertain subsequent threat given that the ‘look’ cue is followed in 50% of the trials by a threatening or neutral stimulus. In the present study, we specifically aimed at examining the pre-stimulus anticipation and stimulus presentation of top-down and bottom-up emotion regulation. Specifically, we focused on the neural activity of the prefrontal cortex and amygdala during pre-stimulus anticipation and stimulus presentation periods. Accumulating evidence indicates that effects of OXT may be moderated by individual difference in baseline trait anxiety levels such that OXT may produce more anxiolytic effect in high trait anxiety individuals (Alvares et al., 2012; Schumacher et al., 2018). In addition, individuals with high trait anxiety show enhanced perceptual sensitivity to threat-related stimuli as well as associated brain activity in amygdala-frontal circuits (Fung et al., 2019). Based on these previous findings we additionally hypothesized that the susceptibility to threatening information and the effects of OXT are modulated by individual variations in trait anxiety.

## Materials and Methods

### Participants

*N* = 88 healthy, right-handed male participants were enrolled in the study. To avoid confounding effects of hormonal changes across the menstrual cycle with OXT administration, the participants were males only. A total of 23 participants were excluded leading to a final sample size of *N* = 65 (randomized double-blind allocation to 35 = OXT, 30 = PLC administration [PLC]; mean age = 20.83 ± 2.60 years). The inclusion and exclusion criteria of the participants are provided in Supplementary material: Participants and CONSORT flow diagram (Figure S1). To further explore the modulatory influence of trait anxiety on OXT effects participants were assigned to a ‘high’ vs. ‘low’ trait anxiety group based on median-split of pre-treatment scores on the State Trait Anxiety Inventory (STAI) (PLC: low anxiety: Mean ± SD = 33.41 ± 3.45, high anxiety: Mean ± SD = 44.38 ± 3.50; OXT: low anxiety: Mean ± SD = 34.19 ± 4.04, high anxiety: Mean ± SD = 45.32 ± 5.31).

The study was pre-registered on the ClinicalTrials.gov database (Identifier: NCT03055546). All participants were free from current or past psychiatric, neurological or other medical disorders (self-report). Participants were instructed to abstain from alcohol and caffeine during the 24 h prior to the experiment. Written informed consent was obtained after a detailed explanation of the study protocol, study procedures had full ethical approval by the local ethics committee and were in accordance with the latest revision of the Declaration of Helsinki.

### Psychometric assessment

Two days before MRI scanning, participants completed the Action Control Scale (ACS-90) by Julius Kuhl (German version: HAKEMP 90, 1990) to examine their self-control ability (Kuhl, 1994). The ACS-90 consists of three subscales: i) action orientation subsequent to failure vs. preoccupation (AOF), ii) prospective and decision-related action orientation vs. hesitation (AOD), iii) action orientation during (successful) performance of activities (intrinsic orientation) vs. volatility (AOP). The AOD subscale was designed to assess prospective and decision-related action orientation, which is related to threat anticipation. Given that the constructs assessed by the AOF and AOP subscales are not relevant for the anticipatory responses, we only focused on the ACS-AOD scale in our analyses. Beck’s Depression Inventory (BDI-II) (Beck et al., 1996) the Autism Spectrum Quotient (AQ) (Baron-Cohen et al., 2001) were administered to control for pre-treatment differences in the levels of depression and autism between treatment groups. Before drug administration and after MRI scanning, participants completed the STAI (Spielberg et al., 1983) and the Positive and Negative Affect Schedule (PANAS) (Watson et al., 1988) scales to evaluate current emotional state. All participants were in the normal range with respect to depression, anxiety and autism scores confirming the self-reported lack of psychiatric conditions in the present sample (Baron-Cohen et al., 2001; Beck et al., 1996).

### Experimental protocol

Figure 1A provides a schematic. overview of the experimental protocol. In a double-blind randomized PLC-controlled between-subject pharmaco-fMRI design, participants self-administered either a single dose of 24 international units (IU) OXT (ingredients: OXT, glycerin, sodium chloride and purified water; Sichuan Meike Pharmaceutical Co. Ltd, Sichuan, China) or PLC nasal spray (produced by Sichuan Meike Pharmaceutical Co. with identical ingredients except for OXT) corresponding to three puffs per nostril. Based on previous pharmaco-kinetic experiments (Paloyelis et al., 2016) treatment was administered 35-40 min before acquisition of the functional MRI time-series.

**Figure 1.**
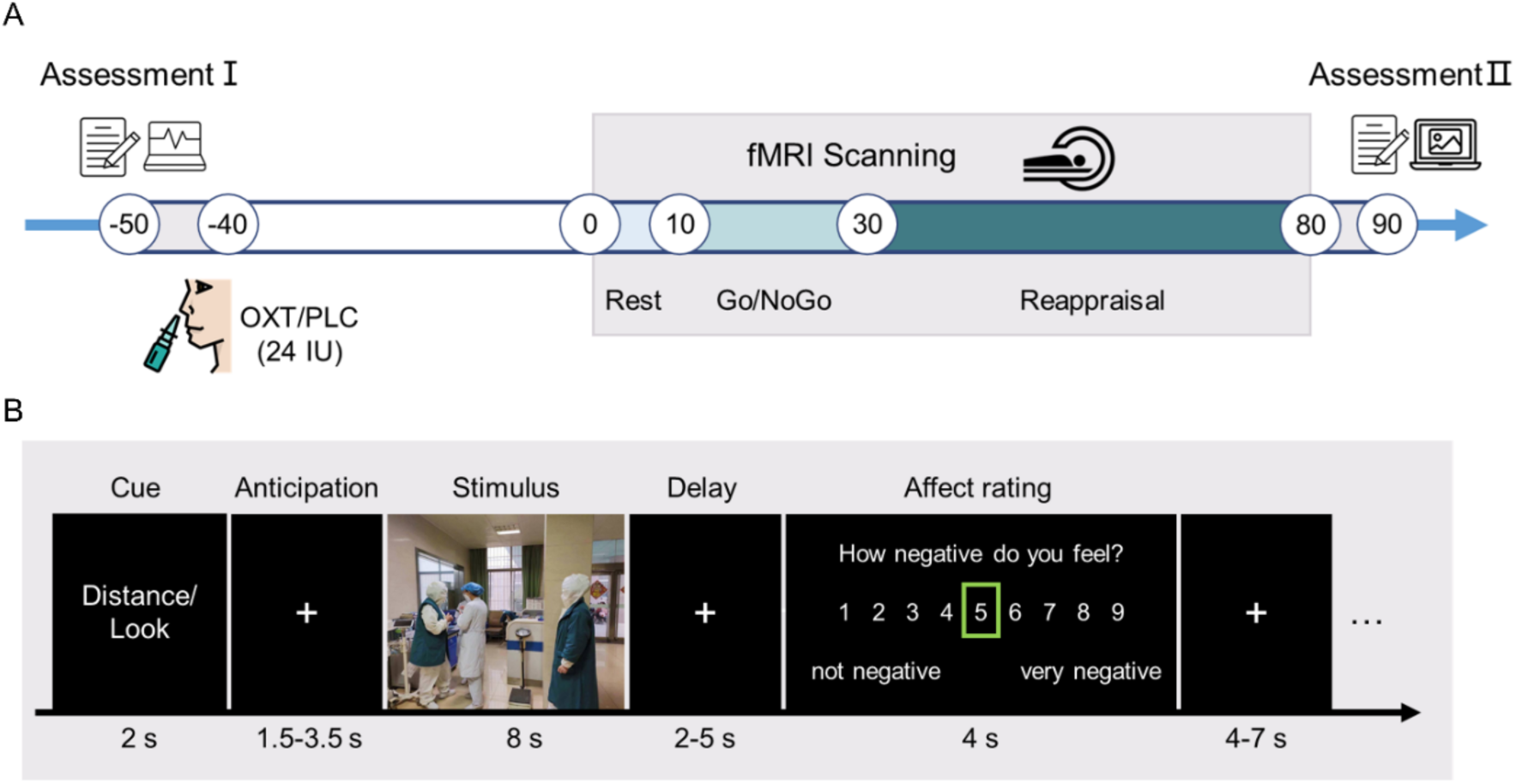
(A) Experimental protocols. (B) Schematic representation and timing of the reappraisal paradigm. For copyright reasons, the picture shown as example is not from the IAPS and permission was obtained.

The subsequent fMRI acquisition included a 7.5 min resting-state scan (Rest), two implicit emotion regulation task runs lasting 4 min 50 s each (emotional Go/NoGo), and six explicit emotion regulation task runs lasting 6 min 46 s each (distancing reappraisal) followed by a surprise memory paradigm outside of the scanner. Results from the implicit emotion regulation and memory paradigm will be reported in separate publications. During the resting-state fMRI acquisition participants were instructed to passively view a fixation cross and keep as motionless as possible. The brain structural data was acquired between the emotional Go/NoGo and distancing reappraisal task, so that there was a five minutes cognitive flushing between two tasks (washout). Head movements were minimized by using comfortable head cushions. In a paper-pencil rating after MRI scanning, the participants answered two questions to evaluate their performance in the reappraisal task: i) indicate how successful they were in regulating their emotion on a scale from 1 (not successful at all) to 7 (very successful), and, ii) can you vividly image the scene/situation depicted in the pictures?

### Stimuli and reappraisal task design

70 minutes after drug administration, participants underwent a distancing reappraisal paradigm with concomitant fMRI acquisition. The design of task is shown in Figure 1B and represents a modified version of prior evaluated event-related cognitive reappraisal paradigms (Koenigsberg et al., 2009; Ochsner et al., 2004; Zimmermann et al., 2017). Each trial began with a 2 s cue (‘Distance’ or ‘Look’), followed by a 1.5-3.5 s anticipatory interval during which a fixation cross was presented. The image was presented for 8 s (aversive or neutral) in a pseudo-random order across participants. Following the stimulus, a fixation cross was presented for a 2–5 s jittered interval followed by rating for 4 s during which participants were asked to rate how negative they felt on a scale of 1-9 (1 = not at all negative, 5 = medium negative, 9 = very negative). The rating was followed by a 4–7 s jittered intertrial interval (ITI).

The task included three trial types: Look Neutral (LookNeu), Look Negative (LookNeg), and Distance Negative (DisNeg). For the ‘Look’ condition, participants were instructed to simply view the image, understand its content, and react naturally without trying to regulate their emotions in any way. For the ‘Distance’ condition, they were instructed to employ cognitive regulation (distancing) in order to view the stimulus from the perspective of a detached and objective observer. The ‘Distance’ cues provided participants a top-down goal to regulate their negative emotion, whereas the ‘Look’ cues simply required participants’ bottom-up emotional response to the subsequent images. Participants were specifically instructed not to close their eyes or look away from the images. The formal experiment did not start until the experimenters confirmed that the participant had fully understood and mastered the distancing reappraisal strategy during a pre-scanning training.

Stimuli for the reappraisal task incorporated 30 neutral and 60 aversive pictures from the International Affective Picture System (IAPS, National Institute of Mental Health Center for Emotion and Attention, University of Florida) database (Lang et al., 1993). Since OXT’s effects were documented as being more pronounced for social stimuli than non-social stimuli, all stimuli depicted threatening social scenes (e.g. corpses, bodily mutilation or assaults) and neutral social situations (e.g. shopping, at work). In a pre-study the pictures were initially rated by an independent sample (n = 37; 18 males) on 9-point scales for the following dimensions: valence (ranging from 1 for unpleasant, to 9 for pleasant), arousal (ranging from 1 for calm, to 9 for aroused). Participants completed six functional runs, each of which contained 15 trials. The last two runs were removed from analyses, since participants’ generally self-reported increased fatigue during the reappraisal task. To objectively confirm across-run fatigue effects (DeLuca et al., 2009), we extracted the parameter estimates during stimuli presentation from the regions involved in visual expertise, including higher visual network, primary visual network and fusiform gyrus. Results revealed reduced activation in these visual regions across the six runs (Supplementary Figure S2) confirming self-reported increasing fatigue over time. Therefore, the final stimuli for all analyses included 20 neutral (‘LookNeu’: valence= 5.33 ± 0.37, arousal = 3.32 ± 0.32; mean ± SD) and 40 negative (20 ‘LookNeg’: valence = 2.33 ± 0.50, arousal = 6.96 ± 0.73; 20 ‘DisNeg’: valence = 2.34 ± 0.54, arousal = 6.96 ± 0.77; mean ± SD) pictures. The valence and arousal of negative pictures were matched between ‘DisNeg’ and ‘LookNeg’ conditions (valence: *P* = 0.995, arousal: *P* = 0.980, two-sample t-tests).

### Behavioral analyses

To identify the main and interaction effects of treatment and trait anxiety on affect rating, affect rating for each participant was subjected to a mixed-effect analysis of variance (ANOVA), with two between-subject factors treatment (OXT, PLC) and anxiety (high, low) and the within-subject factor condition (LookNeu, LookNeg, DisNeg). We also examined the interaction effects of treatment and trait anxiety on reappraisal success using a univariate ANOVA with factors treatment (OXT, PLC) and anxiety (high, low). Reappraisal success was defined as the decrease in reported negative affect rating on DisNeg trials versus LookNeg trials (i.e. LookNeg – DisNeg value) for each participant (similar approach e.g. in Silvers et al., 2015; Zimmermann et al., 2017). Post-hoc pairwise comparisons were corrected for multiple comparisons with Bonferroni corrections. Corresponding analyses were employed using SPSS (version 22.0; IBM SPSS Statistics, Armonk, NY, USA).

### fMRI data acquisition

MRI data was collected on a 3T Siemens Trio scanner (Siemens Magnetom Trio TIM, Erlangen, Germany). High-resolution brain structural data was acquired using a T1-weighted sequence (repetition time [TR] = 1900 ms, echo time [TE] = 2.5 ms, flip angle = 9°, thickness = 1 mm with no gap, field of view [FOV] = 256 mm × 256 mm, image matrix = 256 × 256, 176 sagittal slices, voxel size = 1 × 1 × 1mm) to improve spatial normalization of the functional data. Functional data were acquired using a T2*-weighted Echo Planar Imaging (EPI) sequence with the following parameters: TR/TE = 2000/30ms, flip angle = 90°, acquisition matrix = 64 × 64, voxel size = 3.1 × 3.1 × 3mm, FOV = 200 × 200mm^2^, transverse slices = 33, thickness/gap = 3/0.6 mm.

### fMRI data analysis

fMRI images were preprocessed and analyzed using the standard procedure in SPM12 (http://www.fil.ion.ucl.ac.uk/spm/; Wellcome Trust Centre for Neuroimaging) and DPABI (Version: V4.2_190919; http://rfmri.org/dpabi) (Yan et al., 2016). The first 5 volumes of each functional run were removed to allow for MRI signal equilibrium. The preprocessing steps included slice timing, head motion correction, spatial normalization and smoothing (8 mm full-width at half maximum [FWHM] Gaussian kernel). For each participant and each run, a general linear model (GLM) was constructed by convolving the regressors with the canonical hemodynamic response function (HRF) to estimate evoked blood-oxygen level-dependent (BOLD) activity. The GLM contained regressors indicating the cue (duration = 0 s), anticipation (duration = 1.5-3.5 s, subject-specific time), stimulus (duration = 8 s), rating phase (duration = 4 s), and six motion-correction parameters.

It has been demonstrated that top-down expectation can improve stimulus detection and coding. Expected stimuli can be recognized more rapidly than unexpected ones (Mazzucato et al., 2019; Pinto et al., 2015; Samuelsen et al., 2012; Sussman et al., 2016a). Top-down modulation of visual processing involves visual cortical areas, which is involved with object recognition, as well as parietal and prefrontal cortex, which is involved with visually guided movements and attentional control (Gilbert and Li, 2013). To manifest the mechanism of top-down and bottom-up processes in the distancing reappraisal paradigm, we extracted the time courses from the regions of interest (ROIs) in the primary visual network, higher visual network, fusiform gyrus and visuospatial attention network during stimuli presentation period. The ROIs of visual and visuospatial attention networks were created based on the Stanford Resting-State Network templates (http://findlab.stanford.edu/functional_ROIs.html) (Shirer et al., 2012). The fusiform gyrus mask was created using the Automated Anatomic Labelling (AAL) atlas (Tzourio-Mazoyer et al., 2002) implemented in WFU Pickatlas (Maldjian et al., 2003). The anatomical location and Brodmann areas of each ROI in primary visual, higher visual, fusiform gyrus, and visuospatial attention networks are summarized in Table S1.

To determine the effects of OXT on the anticipation of threat, an ANOVA model was conducted by entering contrast estimates obtained from first-level models into a full factorial model with factors treatment (OXT, PLC) and anticipation (Look, Distance). A separate ANOVA was conducted to examine effects of OXT during the stimulus presentation phase with factors treatment (OXT, PLC) and stimulus type (LookNeu, LookNeg, DisNeg). Results were reported when passing a whole-brain family-wise error (FWE)-corrected extent threshold of *P* < 0.05 at the cluster level or small-volume correction at FWE-corrected threshold of *P* < 0.05 within the anatomically defined bilateral amygdala. The amygdala was chosen based on previous studies demonstrating OXT-induced attenuated amygdala threat reactivity (e.g. Spengler et al., 2017; Striepens et al., 2012). The amygdala mask was created using the AAL atlas (Tzourio-Mazoyer et al., 2002) implemented in WFU Pickatlas (Maldjian et al., 2003). fMRI mean contrast values for subsequent analyses were extracted from a 4-mm radius sphere centered at the peak coordinate of the amygdala cluster.

### Associations between neural and behavioral data

To explore associations between amygdala activity and behavior, Pearson correlations were calculated between the mean beta parameter estimates from the amygdala cluster and behavioral performance (i.e. reappraisal success) across participants. Furthermore, the modulatory influence of individual variations in anxiety (trait anxiety as a measure of bottom-up reactivity) and emotion regulation (ACS-90 scores as a measure of top-down regulatory control) on OXT effects were examined. To this end nonparametric bootstrap tests (10,000 iterations) were conducted on a correlation coefficient to obtain a 95% confidence interval (CI) estimate of Pearson’s r. Between-group correlation differences were tested using Fisher’s z tests. Based on findings from previous studies reporting stronger OXT effects in individuals with higher trait anxiety (Alvares et al., 2012; Schumacher et al., 2018), between-group correlation differences were tested one tailed.

## Results

### Demographics and questionnaires

OXT (n = 35) and PLC (n = 30) groups did not differ in age, levels of depression, mood, anxiety, autistic traits and self-control ability (detailed group characteristics are given in Table 1).

**Table 1.** Demographics and questionnaires

| | PLC<br>($n = 30$) | OXT<br>($n = 35$) | $T$ | $P$ |
| --- | --- | --- | --- | --- |
| **Age (years)** | 20.77 ± 3.05 | 20.89 ± 2.19 | 0.183 | 0.856 |
| **BDI-II** | 6.23 ± 5.33 | 8.03 ± 6.07 | 1.257 | 0.213 |
| **AQ** | 21.53 ± 5.09 | 22.03 ± 4.62 | 0.411 | 0.682 |
| **ACS-AOF** | 5.77 ± 3.21 | 6.03 ± 2.74 | 0.355 | 0.724 |
| **ACS-AOD** | 6.73 ± 2.89 | 6.09 ± 2.67 | -0.938 | 0.352 |
| **ACS-AOP** | 8.80 ± 2.41 | 8.71 ± 1.89 | -0.161 | 0.873 |
| **Pre-Treatment** |  |  |  |  |
| **STAI-state** | 36.67 ± 6.46 | 39.03 ± 6.97 | 1.409 | 0.164 |
| **STAI-trait** | 38.17 ± 6.50 | 40.23 ± 7.33 | 1.191 | 0.238 |
| **PANAS positive** | 23.17 ± 6.54 | 23.49 ± 7.82 | 0.177 | 0.860 |
| **PANAS negative** | 14.70 ± 6.98 | 16.17 ± 6.87 | 0.855 | 0.396 |
| **Post-Scanning** |  |  |  |  |
| **STAI-state** | 36.07 ± 9.76 | 39.97 ± 10.12 | 1.576 | 0.120 |
| **STAI-trait** | 37.53 ± 7.89 | 40.83 ± 9.29 | 1.527 | 0.132 |
| **PANAS positive** | 22.33 ± 6.34 | 19.89 ± 7.45 | -1.413 | 0.162 |
| **PANAS negative** | 13.03 ± 6.57 | 14.23 ± 7.52 | 0.677 | 0.501 |

**Notes:** Data are expressed as mean ± SD (SD: standard deviation). For continuous variables, independent sample $t$ -tests were carried out.

### Behavioral results

We performed a three-way repeated-measures ANOVA, with factors treatment (OXT, PLC), anxiety (high, low) and condition (LookNeu, LookNeg, DisNeg). A significant main effect of condition (*F*[2, 60] = 2300.48, *P* < 0.001, *η*^2^_P_ = 0.987) was obtained, but no main effects of treatment (*P* = 0.505) and trait anxiety (*P* = 0.260). We also did not observe a two-way interaction between treatment and condition (P = 0.420), a two-way interaction between treatment and trait anxiety (*P* = 0.614), or a three-way interaction between the factors (*P* = 0.129). Post hoc tests revealed that: i) negative affect rating was significantly higher during the LookNeg condition than LookNeu condition (*P* < 0.001), confirming a successful induction of moderate negative affect; ii) during the reappraisal (DisNeg) condition significantly less negative affect was reported compared to the LookNeg condition (*P* < 0.001), confirming successful regulation of negative affect in the entire sample (Figure 2A).

**Figure 2.**
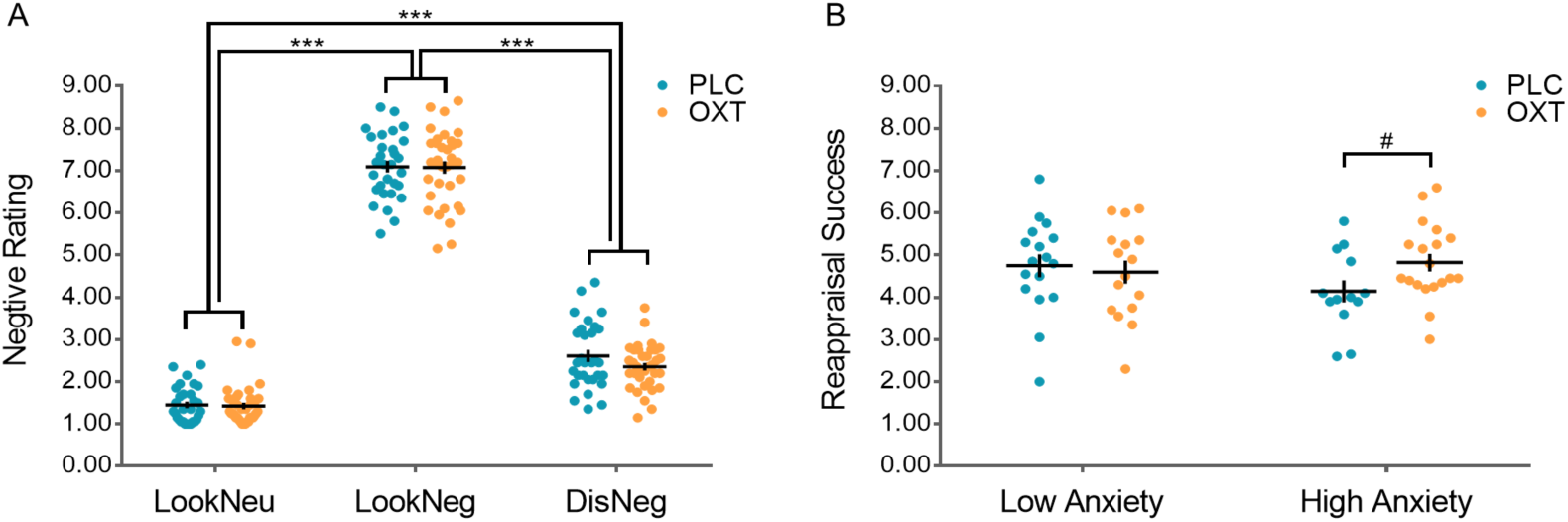
A. Behavioral performance (mean ± standard error) on the emotion regulation task. B. Oxytocin effects on reappraisal success (mean ± standard error) in high and low anxiety groups. 0.05 ≤ #*P* < 0.10; \**P* < 0.05; \*\**P* < 0.01; \*\*\**P* < 0.001.

Based on our hypothesis we explored the interaction effect of treatment and trait anxiety on reappraisal success using a univariate ANOVA with factors treatment (OXT, PLC), anxiety (high, low). There were no significant main effects of treatment (*P* = 0.306) and anxiety (*P* = 0.459), or interaction between treatment and trait anxiety (*P* = 0.108). Based on increasing evidence that OXT may produce more pronounced effects in individuals with high trait anxiety (Alvares et al., 2012; Schumacher et al., 2018) exploratory post hoc simple effects analyses within the anxiety groups were performed. Post hoc Bonferroni-corrected analyses showed a marginally significant improvement in regulation success following OXT in the high anxiety group (reappraisal success: PLC = 4.14 ± 0.94, OXT = 4.82 ± 0.91, *P* = 0.069), see Figure 2B.

### fMRI results

#### Top-down guidance of threat-related perception and attention

In order to investigate the top-down and bottom-up properties of the emotion regulation paradigm, we initially extracted the time courses from the clusters in the primary visual network, higher visual network, fusiform gyrus and visuospatial attention network during stimuli presentation. The time courses confirmed that the ‘Distance’ cue accelerated the onset of stimulus coding, indicating that the visual and attention networks are more rapidly engaged after top-down ‘Distance’ cues compared to bottom-up ‘Look’ cues (see Figure 3).

**Figure 3.**
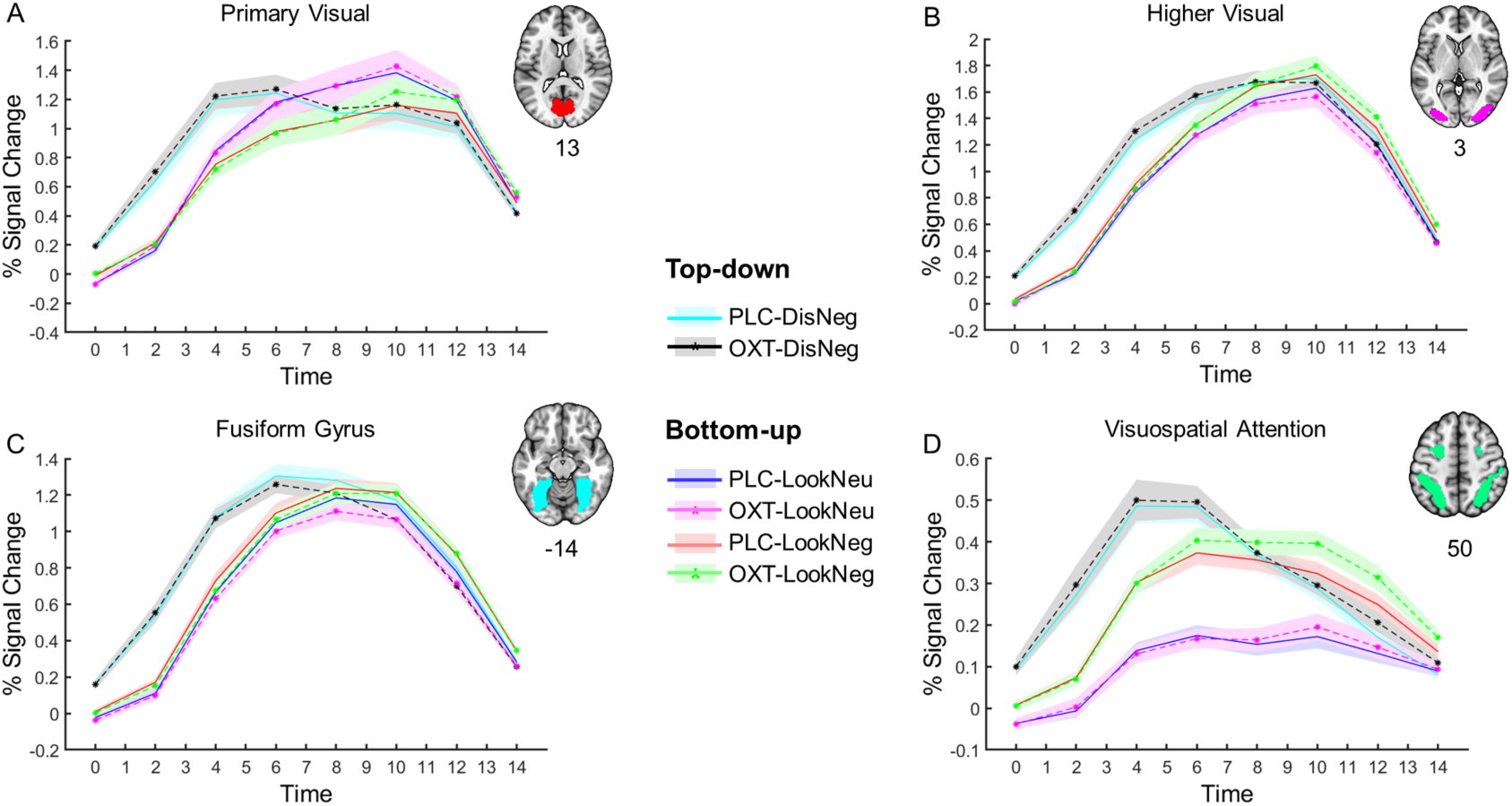
Top-down cue (i.e. ‘Distance’) accelerates stimulus coding in the visual and attention systems. Time courses were extracted from atlas-based regions in the primary visual network (A), higher visual network (B), fusiform gyrus (C), and visuospatial attention network (D).

#### Oxytocin differentially modulated amygdala responses between high and low anxiety during top-down (i.e. distance) and bottom-up (i.e. look) aversive anticipation

The whole-brain full factorial ANOVA model with the factors Treatment (OXT, PLC) and Anticipation (Look, Distance) revealed a significant Treatment × Anticipation interaction effect located in the bilateral posterior insular cortex (left posterior insula/parietal operculum: [-48, -36, 18], *P* = 0.001; right posterior insula/central operculum: [48, -9, 12], *P* = 0.047; cluster-level FWE-corrected) and the right amygdala (right: [33, 0, -24], *P* = 0.001; SV (small volume) FWE-corrected, see Figure S3). Post-hoc examination of the extracted parameter estimates from the regions demonstrating significant interaction effects revealed that OXT significantly attenuated activity in left posterior insula (*t*(63) = - 3.384, *P* = 0.001) and right posterior insula (*t*(63) = -4.200, *P* < 0.001) during distance anticipation in the OXT relative to the PLC group. Moreover, we observed that OXT differentially modulated amygdala activity for ‘Look’ and ‘Distance’ anticipation. Specifically, OXT marginally significantly attenuated the amygdala activity during distance anticipation (*t*(63) = -1.747, *P* = 0.085), but marginally significantly enhanced the amygdala activity during look anticipation (*t*(63) = 1.824, *P* = 0.073). Subsequent extraction of the time-courses from the amygdala region (4-mm radius sphere, centered at [33, 0, -24]) further demonstrated that OXT decreased amygdala activity during the certain threat anticipation (‘Distance’), but increased amygdala activity during uncertain threat anticipation (‘Look’) (Figure 4A and 4B).

**Figure 4.**
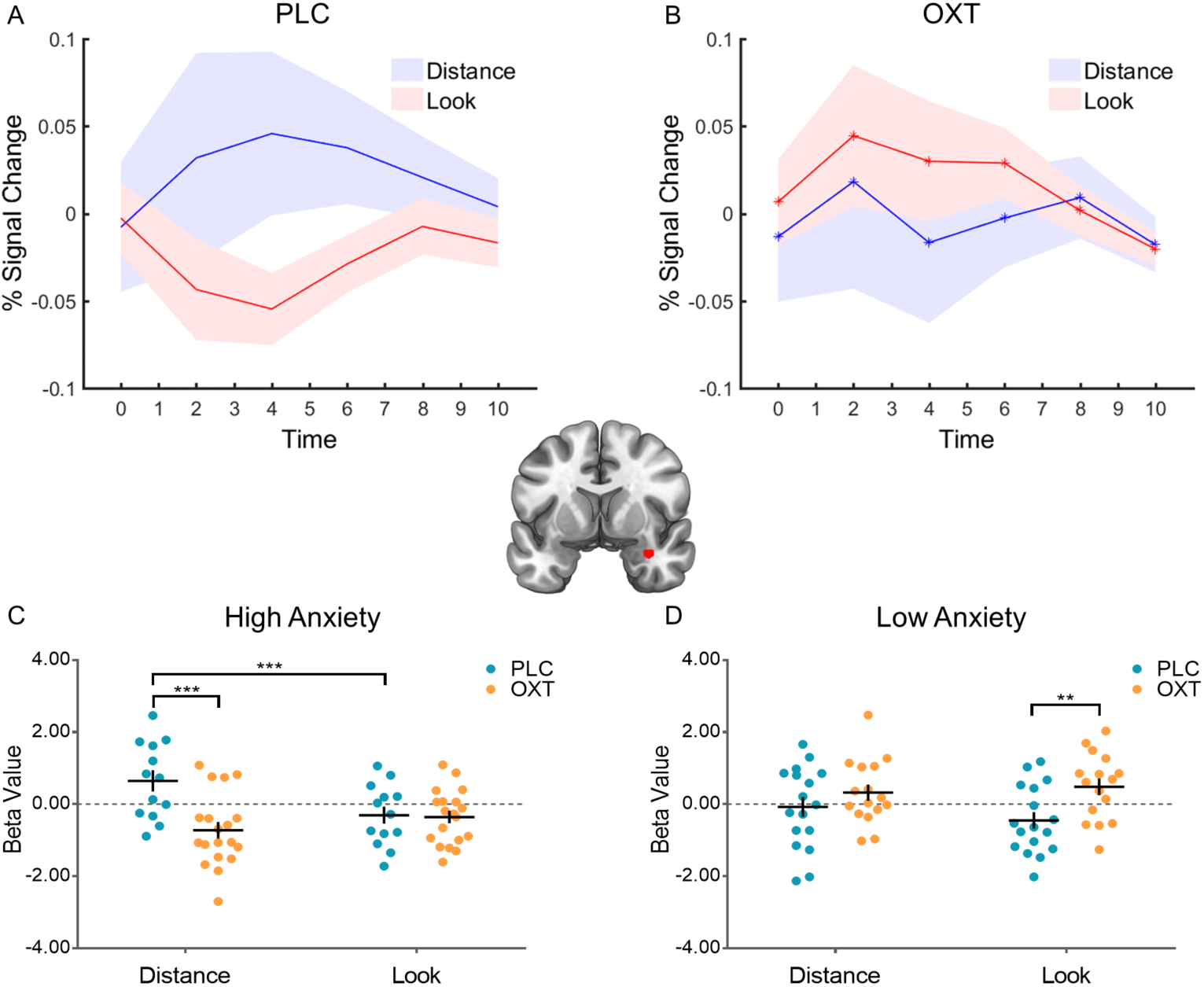
Oxytocin differentially modulated amygdala responses during distance and look aversive anticipation. A. Time course of right amygdala (red cluster) in the (A) PLC group and (B) OXT group. (C) OXT attenuated right amygdala responses during distance anticipation in the high anxiety participants. (D) OXT increased right amygdala responses during look anticipation in the low anxiety participants. \**P* < 0.05; \*\**P* < 0.01; \*\*\**P* < 0.001.

Additional exploratory analyses were conducted to further elucidate the modulatory effect of trait anxiety on the OXT effects. To this end parameter estimates from the amygdala were subjected to a three-way repeated-measures ANOVA with factors Treatment (OXT, PLC), Anxiety (high, low), and Anticipation (Look, Distance). We observed a significant Treatment × Anxiety interaction (*F*[1, 61] = 10.840, *P* = 0.002, *η*^2^_P_ = 0.151) and a marginally significant Treatment × Anxiety × Anticipation interaction effect (*F*[1, 61] = 2.904, *P* = 0.093, *η*^2^_P_ = 0.045). Post hoc Bonferroni-corrected analyses confirmed that the effects of OXT were moderated by individuals’ trait anxiety. Specifically, OXT significantly attenuated anticipatory amygdala activity in the high trait anxiety group (*P* = 0.021), but significantly increased the anticipatory amygdala activity in the low trait anxiety group (*P* = 0.026). An exploratory post hoc 3-way interaction analysis on the extracted parameter estimates from this region furthermore confirmed that OXT significantly attenuated amygdala activity during distance anticipation (*P* < 0.001) in the high trait anxiety group (Figure 4C), but significantly increased amygdala activity during look anticipation (*P* = 0.003) in the low trait anxiety group (Figure 4D).

To explore the Treatment × Anticipation × Anxiety interaction effect in the bilateral posterior insula, parameter estimates from the clusters were subjected to separate three-way mixed ANOVAs with the factors Treatment (OXT, PLC), Anxiety (high, low), and Anticipation (Look, Distance). In the left posterior insula, we observed a significant Treatment × Anxiety interaction (*F*[1, 61] = 5.883, *P* = 0.018, *η*^2^_P_ = 0.088), and a significant Treatment × Anxiety × Anticipation interaction effect (*F*[1, 61] = 8.902, *P* = 0.004, *η*^2^_P_ = 0.127). Post hoc Bonferroni-corrected analyses showed that OXT significantly attenuated the left posterior insula activity during distance anticipation in the high trait anxiety group (*P* < 0.001). In the right posterior insula, we did not observe either main effect or interaction effects involving the factor anxiety.

The whole-brain full factorial ANOVA model with the factors Treatment (OXT, PLC) and Stimulus (LookNeu, LookNeg, DisNeg) did not identify any regions showing either a significant main effect of Treatment or Treatment × Stimulus interaction effects during the stimulus presentation period, suggesting OXT may not modulate explicit cognitive emotion regulation during exposure to threatening stimuli.

#### Associations of anticipatory amygdala activity and subsequent behavior performance

We next examined whether anticipatory amygdala activity was related to subsequent behavior performance, and whether OXT modulated these associations. In general we only observed associations in the PLC group, such that anticipatory amygdala activity following the ‘Distance’ cue showed a significant negative correlation with reappraisal success in the PLC but not in the OXT group (PLC: *r*[28] = -0.430, *P* = 0.018, 95% *CI* = [-0.692; -0.102]; OXT: *r*[33] = 0.021, *P* = 0.904, 95% *CI* = [-0.374; 0.376]; Fisher’s *z*: *z* = 1.84, *P* = 0.033, one tailed; Figure 5A), anticipatory amygdala activity following the ‘Look’ cue showed a significant negative correlation with negative rating of Look trials (LookNeg and LookNeu) in the PLC but not in the OXT group (PLC: *r*[28] = -0.545, *P* = 0.002, 95% *CI* = [-0.758; - 0.260]; OXT: *r*[33] = 0.117, *P* = 0.504, 95% *CI* = [-0.235; 0.429]; Fisher’s *z*: *z* = 2.79, *P* = 0.003, one tailed; Figure 5B). These results indicated that amygdala activity during the pre-stimulus anticipation period was significant associated with individuals’ subsequent emotional reactivity and regulation following PLC, and OXT treatment these associations

**Figure 5.**
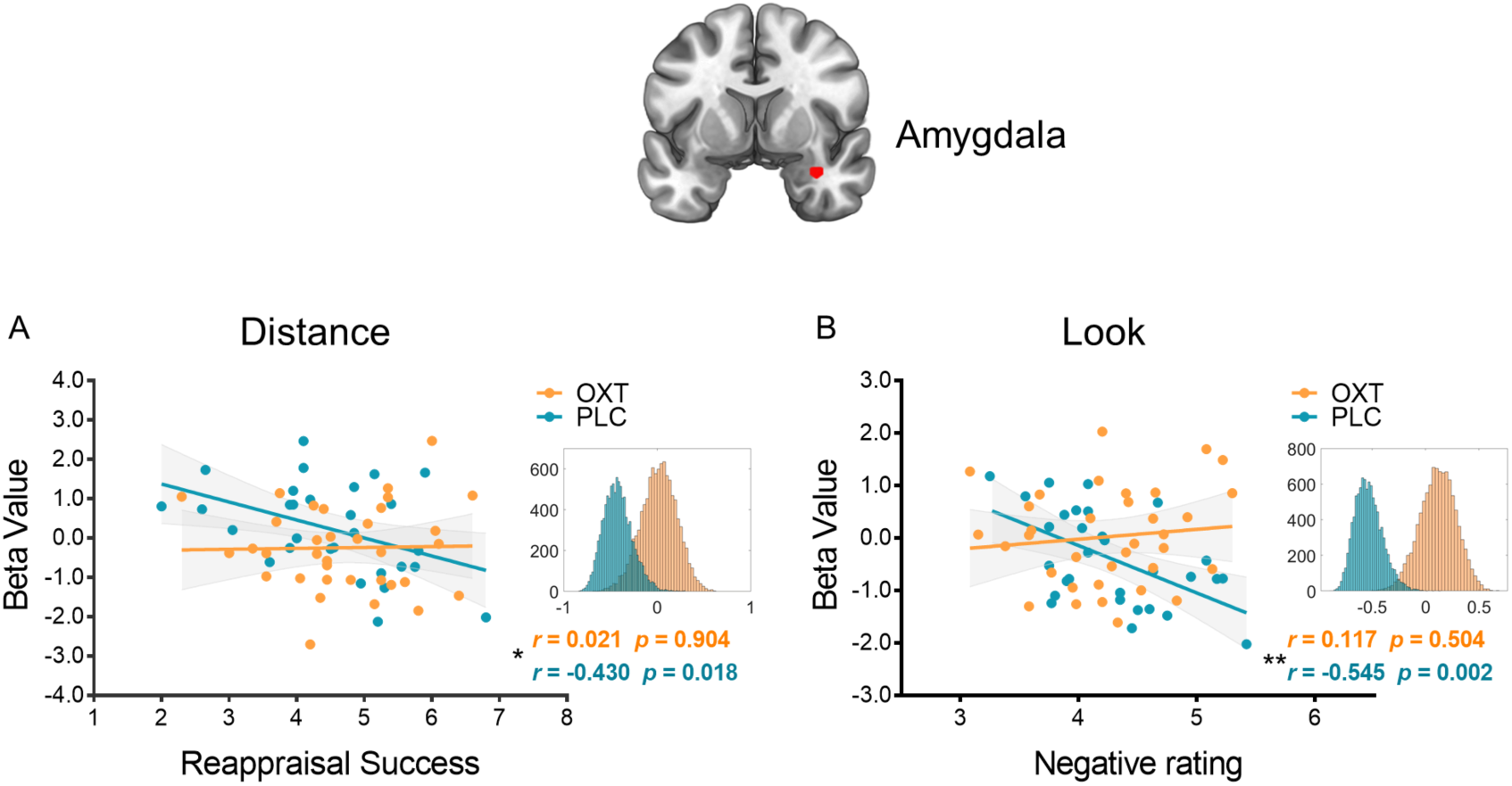
Associations of anticipatory amygdala (red cluster) activity and subsequent behavior performance. (A) A significant negative correlation between anticipatory amygdala activity following the ‘Distance’ cue and reappraisal success in the PLC group; (B) A significant negative correlation between anticipatory amygdala activity following the ‘Look’ cue and negative rating of Look trials in the PLC group. Histograms indicate bootstrapped distribution of correlation coefficients (10,000 iterations). * indicates between-group correlation differences. \**P* < 0.05; \*\**P* < 0.01; \*\*\**P* < 0.001.

#### OXT effects are moderated by Trait anxiety and action control ability

There were no significant pre-treatment differences between treatment groups in trait anxiety (TAI scores) and action control ability (ACS-90 AOD subscale scores; all *Ps* > 0.05, see Table 1). First, we explored associations between TAI and ACS scores and found a significant negative correlation between TAI and ACS-AOD both in the PLC group (*r*[28] = -0.468, *P* = 0.009, 95% *CI* = [-0.724; -0.040]; Figure 6A) and the OXT group (*r*[33] = -0.433, *P* = 0.009, 95% *CI* = [-0.706; -0.130]; Figure 6B).

**Figure 6.**
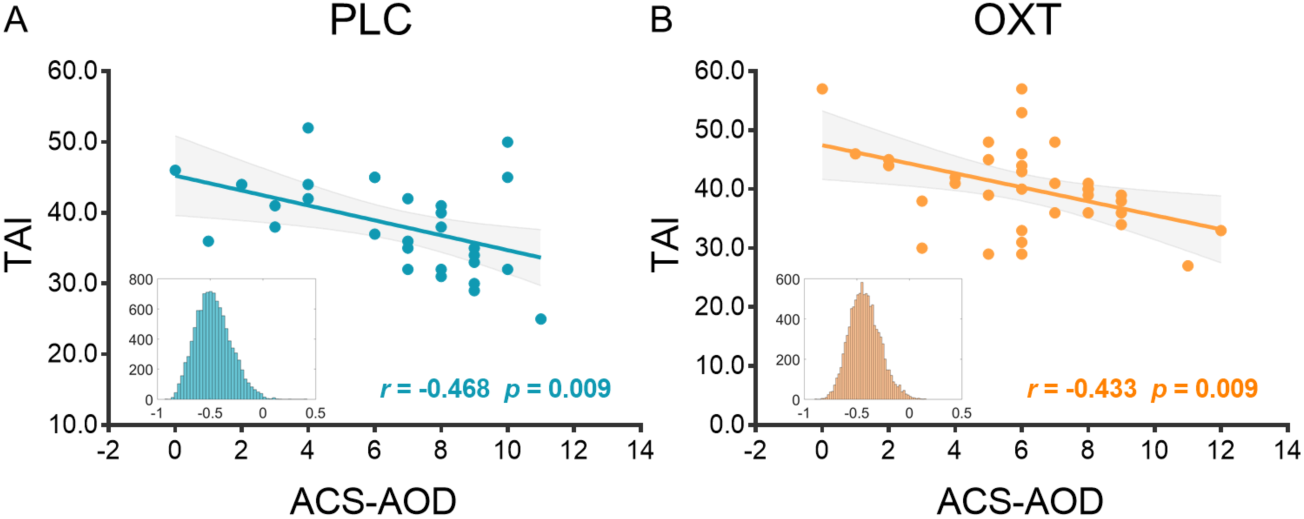
Correlation between STAI and ACS-AOD. A significant negative correlation between TAI and ACS-AOD in the PLC group (A) and the OXT group (B). Histograms indicate bootstrapped distribution of correlation coefficients (10,000 iterations).

We next examined whether anticipatory amygdala activity was related to Trait anxiety and action control ability. In contrast to associations with behavioral indices, the scales only exhibited significant associations in the OXT group: i) anticipatory amygdala activity after ‘Distance’ cue showed significant negative correlation with TAI in the OXT but not in the PLC group (PLC: *r*[28] = 0.246, *P* = 0.190, 95% *CI* = [-0.159; 0.559]; OXT: *r*[33] = -0.567, *P* < 0.001, 95% *CI* = [-0.752; -0.325]; Fisher’s *z*: *z* = 3.42, *P* = 0.0003, one tailed; Figure 7A); ii) anticipatory amygdala activity after ‘Look’ cue showed significant negative correlation with TAI in the OXT but not in the PLC group (PLC: *r*[28] = 0.064, *P* = 0.738, 95% *CI* = [-0.311; 0.374]; OXT: *r*[33] = -0.398, *P* = 0.018, 95% *CI* = [-0.614; -0.125]; Fisher’s *z*: *z* = 1.86, *P* = 0.031, one tailed; Figure 7B); iii) anticipatory amygdala activity after ‘Distance’ cue showed significant positive correlation with ACS-AOD both in the OXT and PLC groups (PLC: *r*[28] = -0.397, *P* = 0.030, 95% *CI* = [-0.690; -0.044]; OXT: *r*[33] = 0.412, *P* = 0.014, 95% *CI* = [0.091; 0.692]; Fisher’s *z*: *z* = 3.28, *P* = 0.0005, one tailed; Fig. 7C); iv) anticipatory amygdala activity after ‘Look’ cue showed significant positive correlation with ACS-AOD both in the OXT and PLC groups (PLC: *r*[28] = -0.408, *P* = 0.025, 95% *CI* = [-0.658; -0.085]; OXT: *r*[33] = 0.514, *P* = 0.002, 95% *CI* = [0.225; 0.732]; Fisher’s *z*: *z* = 3.83, *P* = 0.0001, one tailed; Fig. 7D).

**Figure 7.**
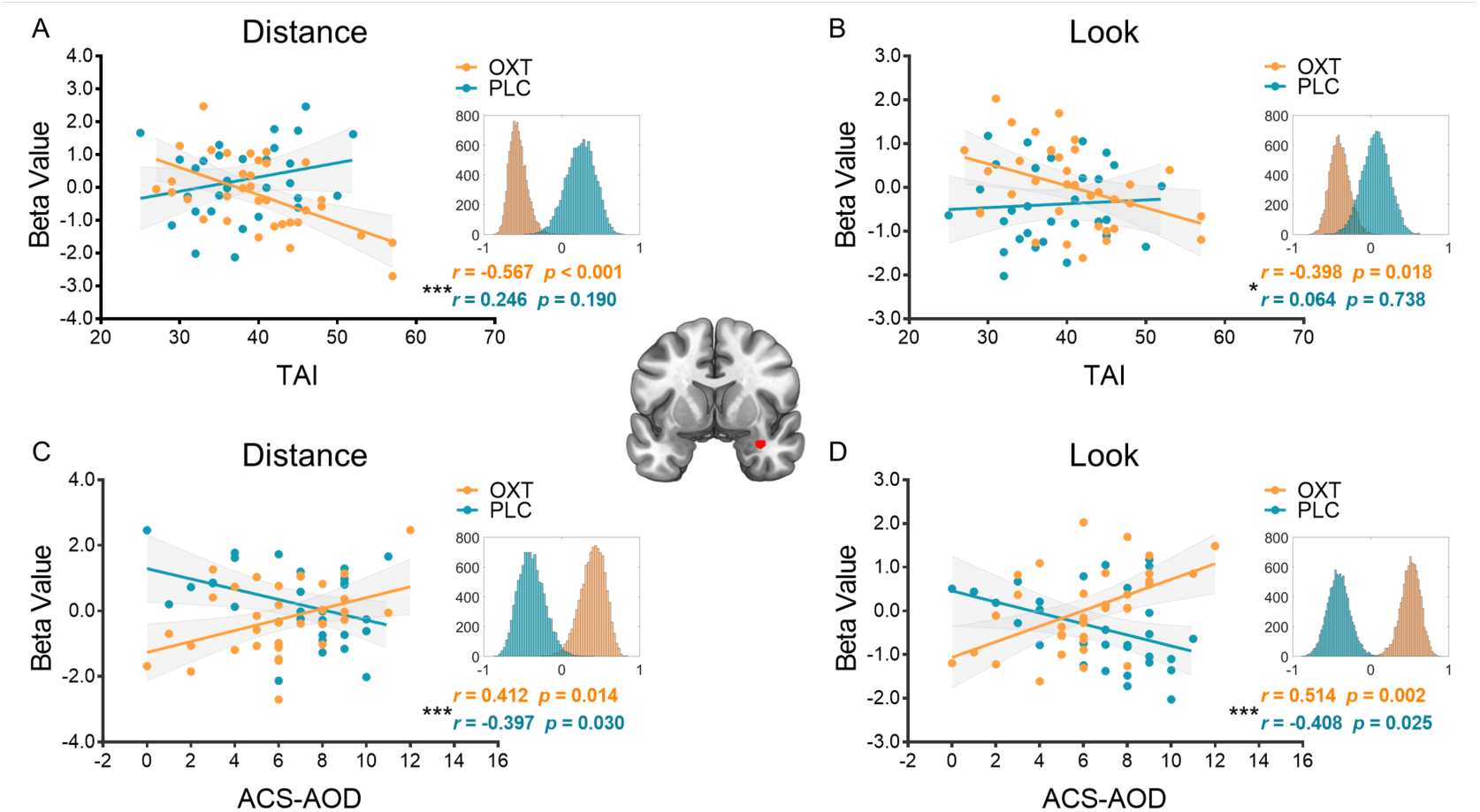
The mean parameter estimates of amygdala (red cluster) in distance anticipation was associated with (A) TAI and (C) ACS-AOD; The mean parameter estimates of amygdala in look anticipation was associated with (B) TAI and (D) ACS-AOD. Histograms indicate bootstrapped distribution of correlation coefficients (10,000 iterations). * indicates between-group correlation differences. \**P* < 0.05; \*\**P* < 0.01; \*\*\**P* < 0.001.

## Discussion

The present pharmacological fMRI study examined the behavioral and neural effects of OXT on top-down and bottom-up emotion generation and regulation by means of a validated emotion regulation paradigm. Our results demonstrate that OXT specifically affected neural activation during the anticipation but not the stimulus presentation phase. Specifically, we observed that OXT attenuated bilateral posterior insular cortex and right amygdala activity during anticipation of top-down regulation of a predictable threat stimulus (‘Distance’), whereas it increased activation in the amygdala during bottom-up anticipation of an unpredictable threat stimulus (‘Look’). Moreover, the effects of OXT on emotion regulation success and anticipatory amygdala activity were modulated by trait anxiety. OXT specifically attenuated amygdala activity during anticipation of top-down regulation and subsequent emotion regulation success in participants with high trait anxiety, whereas it increased anticipatory amygdala activity for bottom-up anticipation in participants with low trait anxiety. Together these findings support a regulatory role of OXT on anxiety-related processing and further emphasize the context- and person-dependent effects of intranasal OXT (Bartz et al., 2011; Olff et al., 2013).

### Top-down guidance of threat-related perception and attention

Previous studies indicate that top-down and bottom-up emotion generation and regulation processes rely on differential neural systems and temporal dynamics (Comte et al., 2016; Ochsner et al., 2009). Relative to stimulus-driven bottom-up processes, top-down processes take into account cues or goals indicating upcoming threat-related stimuli. These anticipatory cues or goals may contribute to perceptual and attentional prioritization of threatening stimuli (Mazzucato et al., 2019; Yoshida and Katz, 2011). To determine the top-down and bottom-up mechanisms during the present emotion regulation task, we extracted the time courses from hub regions of the visual and visuospatial attention networks and observed that top-down anticipatory cues modulated the temporal dynamics of sensory coding and attention control. Specifically, top-down cues accelerated the onset of stimulus coding in visual and visuospatial attention networks, that is, stimuli can be recognized more rapidly following top-down (‘Distance’) cues than bottom-up (‘Look’) cues.

### Oxytocin modulates anticipatory activation in the posterior insula and the amygdala

With respect to OXT effects we found that intranasal OXT attenuated amygdala and posterior insula activation during predictable top-down aversive anticipation and that these effects were mainly driven by treatment effects in participants with high trait anxiety. Specifically OXT increased activation in the amygdala during unpredictable bottom-up aversive anticipation in low trait anxiety individuals. Several previous studies reported OXT attenuated anxiety by reducing amygdala reactivity in response to threatening social stimuli (Eckstein et al., 2015; Grace et al., 2018; Kanat et al., 2015; Spengler et al., 2017), which were primarily interpreted as anxiolytic effects of OXT. On the other hand some studies showed that OXT does not reduce but rather increases threat responses in humans and animals in some contexts (Grillon et al., 2013; Guzmán et al., 2013; Striepens et al., 2012), suggesting that OXT may produce anxiolytic versus anxiogenic effects depending on the specific context as well as individual factors (Bartz et al., 2011; Olff et al., 2013). In contrast to a number of previous studies and our hypothesis, the present study did not reveal effects of OXT on amygdala-threat reactivity or emotion regulation during the stimulus presentation period. Attenuated amygdala threat reactivity following OXT has been previously observed during the presentation of threatening social stimuli (Domes et al., 2007; Eckstein et al., 2015; Kirsch et al., 2005; Labuschagne et al., 2010; Radke et al., 2017). Unlike previous studies, the present paradigm included a preceding cue that allowed participants to infer the probability of a subsequent threat-stimulus and this cue might have shifted the effects of OXT to the anticipation period. The associations between amygdala activation during the anticipation period and level of induction of negative affect and regulation success observed in the placebo group further confirm the OXT’s impact on the anticipation period.

The amygdala plays a critical role in anxiety and fear as well as their regulation across different emotion regulation strategies (Davis, 1992; Etkin et al., 2015; Grupe and Nitschke, 2013; Wager et al., 2008), as well as threat anticipation such that previous human and animal studies reported activation of the amygdala immediately before exposure to an anticipated imminent threat (Davis and Whalen, 2001; Mobbs et al., 2010; Phelps et al., 2001). Clinical studies have repeatedly reported increased amygdala activation during anticipation of aversive stimuli across anxiety disorders, including social anxiety disorder and generalized anxiety disorder (Buff et al., 2017; Figel et al., 2019). The posterior insula/parietal operculum is strongly engaged in interoception, particularly the integration of interoceptive and exteroceptive multimodal sensory information, such as pain, touch, temperature, somato-visceral sensations, auditory processing, and vestibular processing (Craig, 2002; Kurth et al., 2010; Nieuwenhuys, 2012; Simmons et al., 2013; Uddin et al., 2017). Interoceptive information of the current physiological state of the body is progressively integrated with emotional awareness along a posterior to anterior gradient in the insula (Craig, 2002), and interoceptive signaling in the posterior/mid insula has been increasingly recognized as important contributor to emotional experience including anxiety (Tan et al., 2018). Exaggerated anticipation of potential threat represents a key mechanism in anxiety disorders (e.g. Paulus and Stein, 2006) and has been repeatedly associated with exaggerated amygdala and insula activation during the anticipation of threat. For instance, Simmons et al. (2006) found that in anxiety-prone individuals the anticipation of aversive visual stimuli was associated with increased activation in the insular cortex, including the posterior insula, and Tan et al. (2018) reported that posterior/middle insula interoception-related activation was positively associated with trait and state anxiety levels. By using multifaceted anatomical and physiological circuit analyses in mice, Gehrlach et al. (2019) demonstrated a role for the posterior insula in representation of anxiety-related information and described an insula-to-central amygdala pathway which mediates anxiety-related behaviors.

These findings support a key role for the amygdala-posterior insula circuit in anxiety-related processing including threat anticipation (Duval et al., 2015; Shin and Liberzon, 2010; Tovote et al., 2015). Several previous studies reported that OXT modulates amygdala reactivity during threat exposure with increasing evidence for modulatory effects on the posterior insula during anticipation of uncertain and potential painful stimulation (Herpertz et al., 2019) as well as on the functional connectivity of this region with the anterior insula during attentional switching from interoceptive towards external salient cues (Yao et al., 2018). In line with these previous findings, the current results emphasize that OXT the anxiolytic or anxiogenic effects of OXT are neurally mediated by the insula-amygdala circuit.

### Oxytocin differentially modulated amygdala responses in high and low anxiety individuals during top-down (‘Distance’) and bottom-up (‘Look’) aversive anticipation

Exploring the modulatory role of trait anxiety revealed that individual variations in pre-treatment trait anxiety considerably affected the effects of OXT, such that it specifically attenuated amygdala and posterior insula activation during anticipation of top-down regulation and subsequent emotion regulation success in the high anxiety group whereas it increased anticipatory activation in amygdala for bottom-up processing in the low anxiety group. The modulatory role of trait anxiety was further emphasized by correlational analyses that revealed that OXT induced a negative association between trait anxiety and anticipatory amygdala activation for both, top-down and bottom-up anticipation, with visual inspection of the association indicating stronger OXT effects in high anxiety participants during top-down anticipation, yet stronger effects in low anxiety participants during bottom-up anticipation. These findings are in accordance with some prior results (Alvares et al., 2012; Schumacher et al., 2018), suggesting that trait anxiety may be an important modulator of the effects of oxytocin.

Substantial evidence indicates that high anxiety individuals overestimate threat value, which strengthens task-irrelevant stimulus-driven attention but impairs goal-directed attention (Grupe and Nitschke, 2013; Mogg and Bradley, 2016; Sussman et al., 2016a). The imbalance between bottom-up and top-down attention systems may result in emotion dysregulation, excessive anxiety, and threat-related attention bias (Mogg and Bradley, 2018). In contrast, low anxiety individuals may exhibit a lack of adequate arousal and vigilance to cope with unexpected bottom-up threat. In line with this conceptualization individuals with higher trait anxiety demonstrate stronger attentional bias for threat-related stimuli (Broadbent and Broadbent, 1988) allowing faster detection of potential threats in the environment, but deficient cognitive control of task-irrelevant emotional stimuli (Yu et al., 2018). On the neural level individuals with high trait anxiety exhibit increased amygdala activity in response to (Calder et al., 2011; Etkin et al., 2004) as well as during the anticipation of threat stimuli (Calder et al., 2011) and faster escape decisions during threat anticipation in the context of stronger amygdala and insula activation (Fung et al., 2019). This may represent one of the reasons why high anxiety individuals adaptively perform better than low anxiety individuals to identify threat stimuli under anxious conditions (Robinson et al., 2012).

Some evidence suggests that OXT attenuates attentional bias towards threat-related stimuli (Kim et al., 2018; Parr et al., 2013), indicating a role of OXT in attentional control and threat vigilance. In the present study, OXT may thus facilitate top-down goal-directed attention by attenuating amygdala activity in high anxiety participants, while promoting bottom-up attention/vigilance to unexpected threat by enhancing anticipatory amygdala activity in low anxiety participants. The opposite effects of OXT on anticipatory amygdala activation in high versus low anxiety individuals may suggest a baseline anxiety level dependent mechanism via which OXT promotes optimal levels of amygdala activation during the anticipation of an imminent threat. OXT may therefore be a potential intervention to promote an adaptive balance between bottom-up and top-down attention systems depending on individual levels of pre-treatment trait anxiety levels. Given that some disorders are characterized by an imbalance between top-down and bottom-up threat processing, including anxiety disorders (Mogg and Bradley, 2016), schizophrenia (Waters et al., 2012) and autism (Greenaway and Plaisted, 2005), OXT may have a beneficial therapeutic effect in these disorders.

The cognitive processes of distancing reappraisal during the anticipation period mainly involves attentional control, whereas the stimulus presentation period additionally involves self-projection, affective self-reflection, and cognitive control (Koenigsberg et al., 2010; Ochsner et al., 2012; Powers and LaBar, 2019). Combined with findings from our prior resting state study (Xin et al., 2018), it may be hypothesized that OXT may modulating pre-stimulus attentional control of threatening social information. With respect to behavior performance during emotion regulation, no effects of OXT per se were observed, although the high anxiety group exhibited a marginal significant improvement of reappraisal success following OXT, suggesting that OXT may have the potential to improve cognitive emotion regulation in high anxiety individuals. In addition, we predicted that OXT would increase prefrontal cortex (PFC) activity during distancing reappraisal, and thus enhance explicit emotion regulation. However, we did not observe OXT effects on PFC during emotion regulation. In accordance with prior findings, we observed that distancing reappraisal mainly recruited the default network (Koenigsberg et al., 2010; Koenigsberg et al., 2009). The default network, comprising the precuneus/posterior cingulate cortex, anterior medial prefrontal cortex, medial temporal and inferior parietal regions, plays key role in internally oriented and self-referential mental processes (Buckner and DiNicola, 2019; Raichle et al., 2001). This finding again supports the idea that distancing reappraisal involves internally-oriented attention, which recruits the default network. The primary engagement of the posterior regions by the current paradigm may have contributed to the absence of OXT effects on the PFC in the present study.

### Associations of anticipatory amygdala activity and subsequent behavior performance

Anticipation is a universal preparatory response to an upcoming or uncertain event and previous studies reported that pre-stimulus anticipatory neural activity predicts the subsequent emotion regulation success. For instance, pre-stimulus functional connectivity between key areas of pain relates to the susceptibility to pain (Ploner et al., 2010). Moreover, Denny et al. (2014) found that anticipatory brain activity predicts the success or failure of subsequent emotion regulation. In the present study, anticipatory amygdala activity was associated with subsequent emotion regulation performance only in the PLC group. Specifically, i) lower anticipatory amygdala activity after the ‘Distance’ cue associated with higher reappraisal success, and, ii) higher anticipatory amygdala activity after the ‘Look’ cue associated with lower negative affect ratings. In sum, higher top-down anticipatory amygdala activity was associated with improved explicit emotion regulation performance, whereas higher bottom-up anticipatory amygdala activity with lower induction of negative affect by the subsequent stimulus. The associations between pre-stimulus amygdala activity and subsequent performance were not observed following OXT, indicating that OXT uncoupled the association between anticipatory amygdala activation and subsequent emotional experience and emotion regulation.

### Exploratory analysis of associations between action control ability and anticipatory amygdala activity under OXT and PLC

We additionally explored associations between individual variations in action control ability (as assessed by the ACS-90 scale) and anticipatory amygdala activity. While no associations between anticipatory amygdala activity and trait anxiety levels were observed in the PLC group, we observed significant negative associations with action control ability in this group but significant positive associations in the OXT group. Visual inspection of the scatterplots suggests that OXT primarily decreased amygdala activity in individuals with a low action control ability during the top-down (‘Distance’) anticipatory period, whereas increased amygdala activity of the individuals with high action control ability during the bottom-up (‘Look’) anticipatory period. These findings indicate that the effects of OXT are not only modulated by ‘emotional reactivity’ traits (anxiety) but also by emotion regulation relevant traits (action control).

### Cognitive reappraisal from the perspective of large-scale brain networks

At the neural level, emotion regulation requires the concerted engagement of districted brain systems beyond prefrontal regions such as dorsolateral PFC, dorsomedial PFC, and ventromedial PFC (Etkin et al., 2015; Ochsner et al., 2012). For instance, in the present paradigm successful emotion regulation via distancing was accompanied by increased activation in multiple brain systems, such as the posterior default network, dorsal attention network, frontoparietal control network and salience network (see Supplementary material). Research on interacting large-scale networks (Bassett and Sporns, 2017; Cocchi et al., 2013; Cohen and D’Esposito, 2016; Shine and Poldrack, 2018; Sporns, 2018) that support successful emotion regulation may thus provide novel insights into the underlying brain networks.

### Limitations and conclusions

The present study focused on the effects of OXT in men and in the context of increasing evidence for sexual dimorphic effects of OXT on amygdala and insula responses (e.g. Gao et al., 2016; Ma et al., 2018) the generalization of the findings to women remains to be determined in future studies. The present study employed a cognitive emotion regulation paradigm to examine the modulatory effect of intranasal OXT on bottom-up and top-down emotion generation and regulation during the anticipation and presentation of threat stimuli. We found that OXT attenuated bilateral posterior insular cortex and right amygdala activation during anticipation of top-down regulation of predictable threat stimuli in participants with high trait anxiety, whereas it increased activation in the amygdala during anticipation of bottom-up unpredictable threat stimuli in participants with low trait anxiety. Collectively, these findings support a regulatory role of OXT on threat anticipation and further emphasize the context- and person-dependent effects of intranasal OXT.

## Acknowledgments

This work was supported by the National Key Research and Development Program of China (Grant No. 2018YFA0701400), National Natural Science Foundation of China (NSFC, No 91632117; 31530032); Fundamental Research Funds for Central Universities (ZYGX2015Z002), and, Science, Innovation and Technology Department of the Sichuan Province (2018JY0001).

## Authors contributions

Author contributions are as follows: design, F.X. and B.B.; data collection, F.X., X.Z., Z.Z., X.Y., Q.W. and Y.G.; data analysis, F.X., X.Z., and D.D.; interpretation of data and drafting of the manuscript F.X., B.B., A.C. and K.M.K. We would like to thank Feng Zhou and Xiaole Ma for help with data collection.

