## Supplementary Information for "Oxytocin differentially modulates amygdala responses during top-down and bottom-up aversive anticipation"

### Supplementary material

**Xin et al.,**

#### Participants

N = 88 healthy, right-handed male participants were enrolled in the study. A total of 23 participants were excluded leading to a final sample size of N = 65. Three participants were excluded due to incomplete data. The present study involved emotion perception and emotion regulation, which has been demonstrated to be influenced by individuals' depressive and autistic traits (Poljac et al., 2013; Rive et al., 2013). Four participants with high depressive and autistic traits were therefore excluded from analyses (BDI-II > 28; AQ > 30, in ref. Liu, 2008). In order to match the trait anxiety scores between OXT and PLC groups, three participants with high trait anxiety (TAI > 60) in the OXT group were excluded. Four participants were excluded due to excessive head motion (>3 mm translation, >3° rotation). One participant whose mean negative rating of reappraisal stimuli (i.e. DisNeg) was beyond three standard deviations had to be excluded. Six participants whose mean negative ratings of neutral stimuli (i.e. LookNeu) were larger than 3 had to be excluded. Two participants whose accuracy of surprise memory test were beyond three standard deviations had to be excluded. Figure S1 showed the CONSORT flow diagram.

CONSORT
TRANSPARENT REPORTING of TRIALS

### CONSORT 2010 Flow Diagram

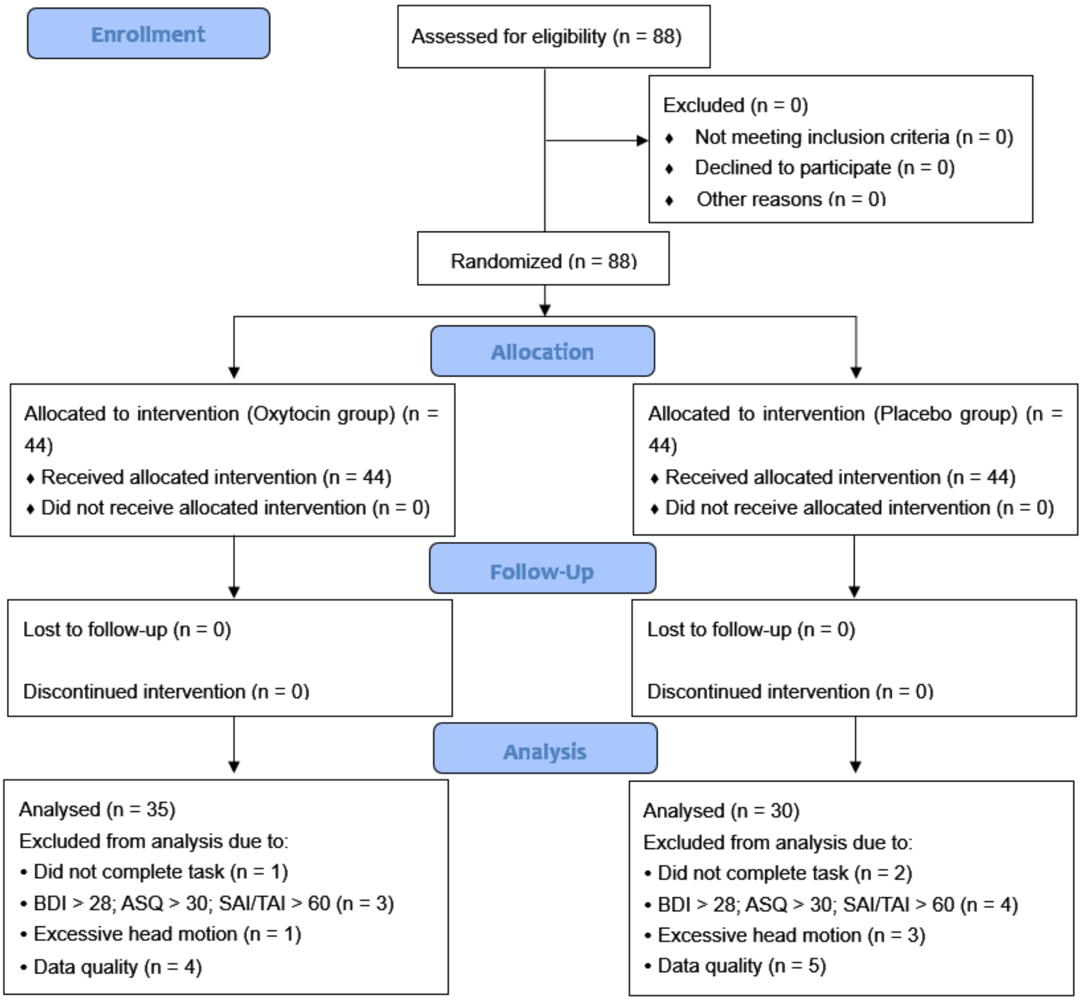


**Figure S1**. The CONSORT flow diagram.

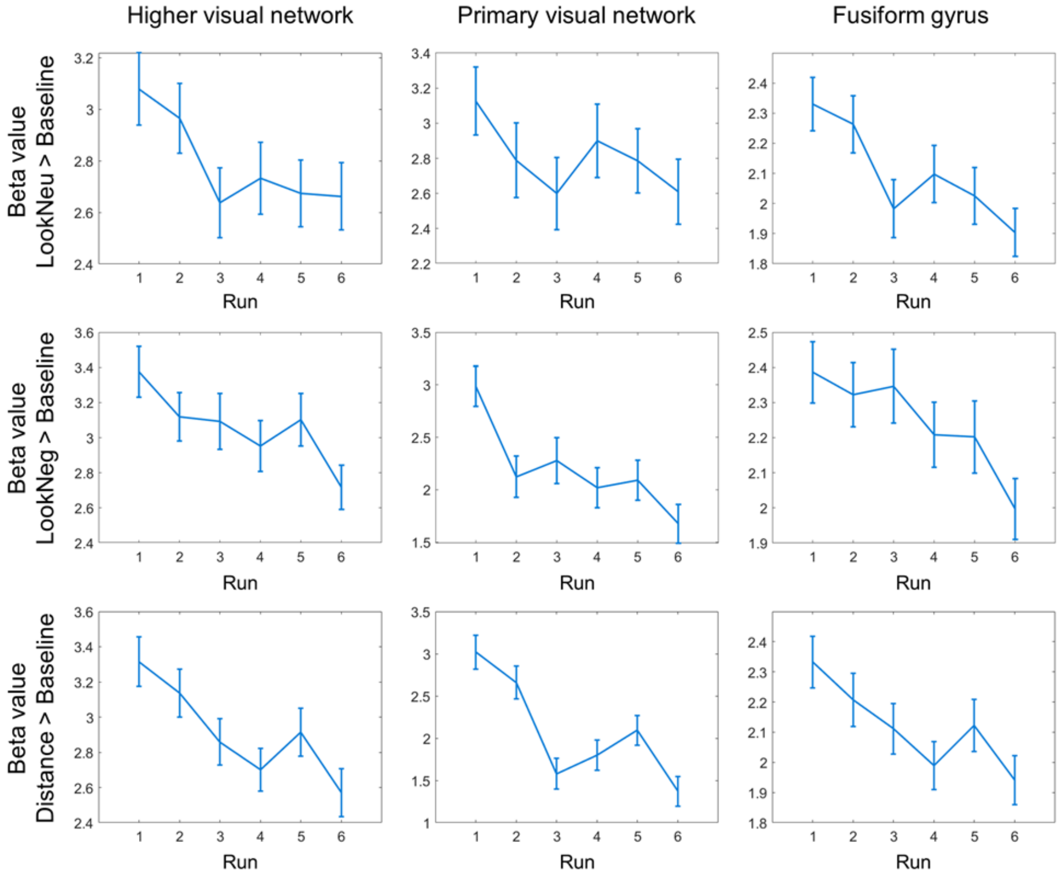


**Figure S2.** Across-run fatigue. Reduced activation in the higher visual network, primary visual network and fusiform gyrus during the stimuli presentation across the six runs.

**Table S1.** The anatomical location and Brodmann areas of each ROI in primary visual network, higher visual network, fusiform gyrus, and visuospatial attention network.

| Network | Anatomical Location of Functional ROIs | Brodmann Areas |
|---|---|---|
| Primary Visual Network | Calcarine Sulcus | 17 |
| Higher Visual Network | Left Middle Occipital Gyrus, Superior Occipital Gyrus | 18, 19, 17 |
| | Right Middle Occipital Gyrus, Superior Occipital Gyrus | 17, 18, 19 |
| Visuospatial Attention Network | Left Middle Frontal Gyrus, Superior Frontal Cortex, Precentral Gyrus | 6 |
| | Left Inferior Parietal Sulcus | 2, 40, 7 |
| | Left Frontal Operculum, Inferior Frontal Gyrus | 44, 48, 45 |
| | Left Inferior Temporal Gyrus | 37 |
| | Right Middle Frontal Gyrus | 6 |
| | Right Inferior Parietal Lobule | 2, 40, 7 |
| | Right Frontal Operculum, Inferior Frontal Gyrus | 44, 48 |
| | Right Middle Temporal Gyrus | 37 |
| Fusiform Gyrus | Left Fusiform | 37 |
| | Right Fusiform | 37 |

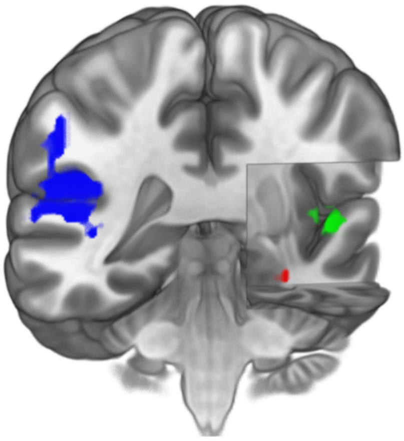

**Figure S3.** A significant interaction between treatment and anticipation was observed in posterior insula (left: [-48, -36, 18], blue; right: [48, -9, 12], green) and amygdala (right: [33, 0, -24], red).

**General activation patterns engaged during the paradigm**

Examining the DisNeg vs. LookNeg contrast during stimuli presentation revealed widespread activity in the posterior default network and frontoparietal control network (see Figure S4A and Table S2). The LookNeg vs. DisNeg contrast during stimuli presentation showed activity in visual network, fusiform gyrus and part of dorsal attention network (see Figure S4B and Table S3). The DisNeg vs. LookNeg contrast during pre-stimulus anticipation period revealed that activity in the posterior default network and parts of frontoparietal control network diminished during reappraisal (see Figure S6 and Table S4). The LookNeg vs. DisNeg contrast during pre-stimulus anticipation revealed no significant differences. Together, these results are consistent with prior studies (Koenigsberg et al., 2009; Xie et al., 2016), indicating that our distancing reappraisal paradigm engaged the previously reported emotion regulation networks. We additionally extracted the time courses from the clusters in the visual and attention networks in the LookNeg vs. DisNeg contrast during stimuli presentation. Similarly, time courses showed that 'Distance' cue accelerates onset of stimulus coding, that is visual and attention networks can be activated more rapidly after top-down 'Distance' cues than bottom-up 'Look' cues (see Figure S5).

**Table S2**. DisNeg vs. LookNeg contrast during stimuli presentation.

| Region | H. | Abbr. | MNI-coordinates | | | Cluster Size | Peak *T* |
|---|---|---|---|---|---|---|---|
| | | | x | y | z | | |
| Stimuli presentation：DisNeg-LookNeg (OXT+PLC) | | | | | | | |
| Precuneus | L | PCu | -9 | -57 | 33 | 832 | 11.677 |
| Cuneus | R | Cun | 9 | -90 | 21 | | 8.428 |
| Cuneus | L | Cun | -3 | -93 | 15 | | 7.902 |
| Angular gyrus | L | AnG | -45 | -66 | 39 | 515 | 9.863 |
| Angular gyrus | L | AnG | -48 | -63 | 30 | | 9.572 |
| Superior temporal gyrus | R | STG | 69 | -36 | 3 | 152 | 6.974 |
| Middle temporal gyrus | R | MTG | 51 | -33 | 0 | | 6.638 |
| Middle temporal gyrus | L | MTG | -63 | -36 | -3 | 99 | 6.964 |
| Middle frontal gyrus | L | MFG | -39 | 6 | 54 | 88 | 6.181 |
| Middle frontal gyrus | L | MFG | -39 | 15 | 45 | | 5.525 |
| Middle temporal gyrus | L | MTG | -54 | -3 | -24 | 78 | 5.978 |
| Middle temporal gyrus | L | MTG | -60 | -18 | -21 | | 5.239 |
| Superior frontal gyrus | L | SFG | -12 | 18 | 63 | 10 | 5.473 |
| Middle cingulate gyrus | L | MCgG | 0 | -21 | 36 | 3 | 5.138 |
| Superior frontal gyrus | R | SFG | 15 | 18 | 60 | 3 | 5.126 |
| Middle temporal gyrus | R | MTG | 66 | -9 | -24 | 3 | 5.091 |
| Superior frontal gyrus | L | SFG | -15 | 51 | 33 | 2 | 4.985 |

H, hemisphere; MNI, Montreal Neurological Institute; L, left; R, right. Whole-brain FWE corrected at peak-level, P < 0.05.

**Table S3**. LookNeg vs. DisNeg contrast during stimuli presentation.

| Region | H. | Abbr. | MNI-coordinates x | y | z | Cluster Size | Peak *T* |
|---|---|---|---|---|---|---|---|
| Stimuli presentation：LookNeg – DisNeg (OXT+PLC) | | | | | | | |
| Lingual gyrus | R | LiG | 6 | -66 | 3 | 1011 | 10.505 |
| Fusiform gyrus | R | FuG | 30 | -45 | -9 | | 10.179 |
| Lingual gyrus | L | LiG | -3 | -66 | 3 | | 9.918 |
| Middle occipital gyrus | L | MOG | -33 | -84 | 24 | 172 | 9.858 |
| Fusiform gyrus | L | FuG | -27 | -45 | -12 | 532 | 9.005 |
| Inferior occipital gyrus | L | IOG | -24 | -93 | -6 | | 8.117 |
| Fusiform gyrus | L | FuG | -30 | -60 | -9 | | 7.943 |
| Postcentral gyrus | L | PoG | -45 | -30 | 48 | 633 | 8.589 |
| Postcentral gyrus | L | PoG | -54 | -24 | 27 | | 8.409 |
| Precentral gyrus | L | PrG | -39 | -18 | 63 | | 7.965 |
| Middle occipital gyrus | R | MOG | 39 | -78 | 30 | 375 | 8.109 |
| Superior parietal lobule | R | SPL | 30 | -63 | 42 | | 6.290 |
| Superior parietal lobule | R | SPL | 30 | -54 | 48 | | 5.874 |
| Supramarginal gyrus | R | SMG | 51 | -27 | 45 | 222 | 6.708 |
| Supramarginal gyrus | R | SMG | 60 | -18 | 30 | | 5.724 |
| Precentral gyrus | L | PrG | -48 | 3 | 30 | 41 | 6.315 |
| Precentral gyrus | R | PrG | 51 | 9 | 30 | 44 | 6.259 |
| Inferior temporal gyrus | R | ITG | 54 | -48 | -12 | 42 | 6.251 |
| Amygdala | R | AmG | 21 | 0 | -18 | 11 | 5.834 |
| Superior parietal lobule | L | SPL | -21 | -63 | 45 | 31 | 5.798 |
| Posterior insula | L | PIns | -36 | -3 | -9 | 18 | 5.549 |
| Middle frontal gyrus | R | MFG | 27 | 3 | 57 | 2 | 5.002 |
| Precuneus | R | PCu | 24 | -57 | 18 | 1 | 4.965 |

H, hemisphere; MNI, Montreal Neurological Institute; L, left; R, right. Whole-brain FWE corrected at peak-level, $P < 0.05$.

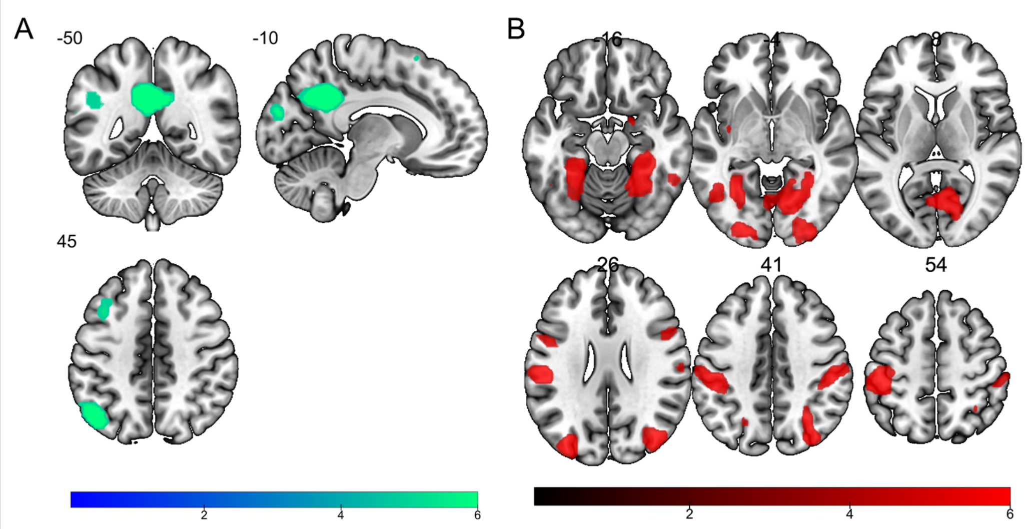


**Figure S4**. (A) DisNeg vs. LookNeg contrast and (B) LookNeg vs. DisNeg contrast during stimuli presentation.

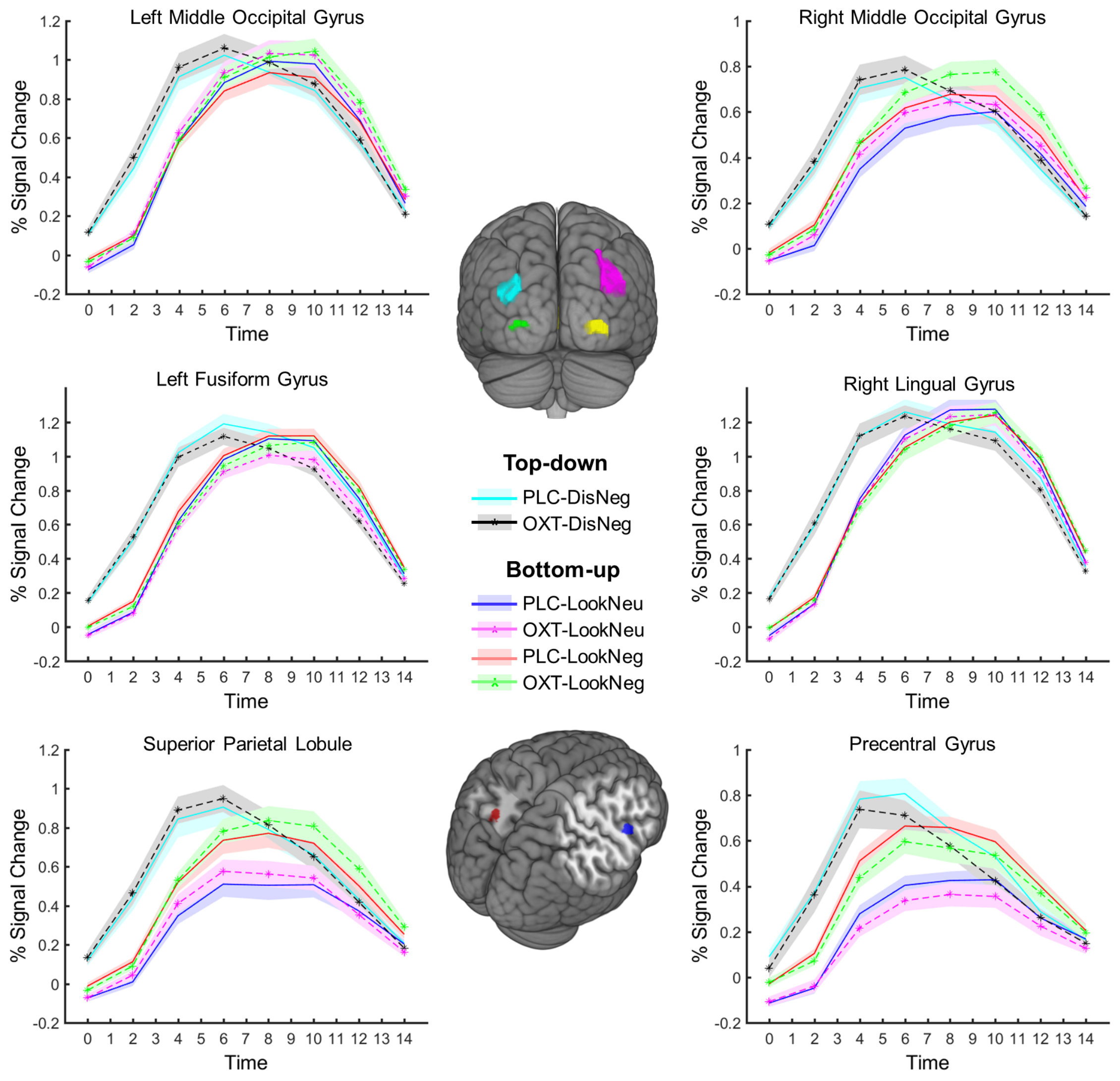


**Figure S5.** Top-down cue (i.e. 'Distance') accelerates stimulus coding in the visual perception and sensory. Time courses reflect that the perception cortex can be activated more rapidly after 'Distance' cue than 'Look' cue. Time courses were extracted from the clusters in visual and attention networks in the LookNeg vs. DisNeg contrast during stimuli presentation. All regions survived peak-level FWE correction, P < 0.05, see Table S3. Cyan: Left middle occipital gyrus, [-33, -84, 24]; Violet: Right Middle Occipital Gyrus, [39, -78, 30]; Green: Left fusiform gyrus, [-27, -45, -12]; Yellow: Right lingual gyrus, [6, -66, 3]; Red: Superior Parietal Lobule, [-21, -63, 45]; Blue: Precentral Gyrus, [51, 9, 30].

**Table S4**. DisNeg vs. LookNeg contrast during pre-stimuli anticipation.

| Region | H. | Abbr. | MNI-coordinates x | y | z | Cluster Size | Peak *T* |
|---|---|---|---|---|---|---|---|
| Pre-stimuli anticipation: DisNeg - LookNeg (OXT+PLC) | | | | | | | |
| Precentral gyrus | L | PrG | -51 | 6 | 39 | 135 | 7.660 |
| Angular gyrus | L | AnG | -45 | -63 | 15 | 296 | 7.405 |
| Occipital fusiform gyrus | L | OFuG | -36 | -72 | -12 | | 6.768 |
| Middle temporal gyrus | L | MTG | -57 | -45 | -12 | | 6.308 |
| Angular gyrus | L | AnG | -39 | -66 | 42 | 233 | 7.226 |
| Angular gyrus | L | AnG | -39 | -54 | 45 | | 6.241 |
| Inferior occipital gyrus | R | IOG | 42 | -75 | 9 | 116 | 7.011 |
| Middle occipital gyrus | R | MOG | 33 | -78 | 18 | | 6.495 |
| Precuneus | L | PCu | -3 | -75 | 39 | 300 | 6.763 |
| Precuneus | L | PCu | -3 | -66 | 36 | | 6.185 |
| Calcarine cortex | L | Calc | -12 | -72 | 9 | | 6.096 |
| Middle cingulate gyrus | L | MCgG | -6 | -24 | 42 | 48 | 6.726 |
| Posterior cingulate gyrus | L | PCgG | -3 | -33 | 36 | | 5.527 |
| Occipital fusiform gyrus | R | OFuG | 36 | -63 | -12 | 235 | 6.581 |
| Occipital fusiform gyrus | R | OFuG | 30 | -75 | -12 | | 6.381 |
| Occipital fusiform gyrus | R | OFuG | 30 | -63 | -6 | | 6.310 |
| Supramarginal gyrus | R | SMG | 51 | -27 | 42 | 120 | 6.454 |
| Superior parietal lobule | R | SPL | 39 | -45 | 57 | | 5.754 |
| Supramarginal gyrus | R | SMG | 60 | -21 | 27 | | 5.488 |
| Precentral gyrus | R | PrG | 48 | 9 | 27 | 124 | 6.395 |
| Precentral gyrus | R | PrG | 51 | 9 | 39 | | 6.268 |
| Middle frontal gyrus | L | MFG | -33 | 3 | 66 | 66 | 6.304 |
| Middle frontal gyrus | L | MFG | -39 | 9 | 57 | | 5.871 |
| Thalamus proper | L | TP | -3 | -15 | 12 | 15 | 5.786 |
| Posterior cingulate gyrus | L | PCgG | -6 | -45 | 9 | 13 | 5.737 |
| Precentral gyrus | R | PrG | 45 | -6 | 54 | 18 | 5.592 |
| Superior frontal gyrus | L | SFG | -15 | 12 | 63 | 7 | 5.578 |
| Triangular part of the inferior frontal gyrus | L | TrIFG | -48 | 36 | 3 | 6 | 5.453 |
| Precentral gyrus | L | PrG | -24 | -21 | 72 | 4 | 5.360 |
| Superior frontal gyrus | L | SFG | -15 | 6 | 72 | 2 | 5.268 |
| Precentral gyrus | R | PrG | 39 | -21 | 69 | 3 | 5.245 |
| Lingual gyrus | L | LiG | -3 | -75 | -3 | 5 | 5.189 |
| Posterior cingulate gyrus | R | PCgG | 6 | -42 | 0 | 8 | 5.150 |
| Posterior cingulate gyrus | R | PCgG | 9 | -42 | 9 | | 5.149 |
| Superior frontal gyrus | L | SFG | -27 | 15 | 63 | 3 | 5.115 |

H, hemisphere; MNI, Montreal Neurological Institute; L, left; R, right. Whole-brain FWE corrected at peak-level, $P < 0.05$.

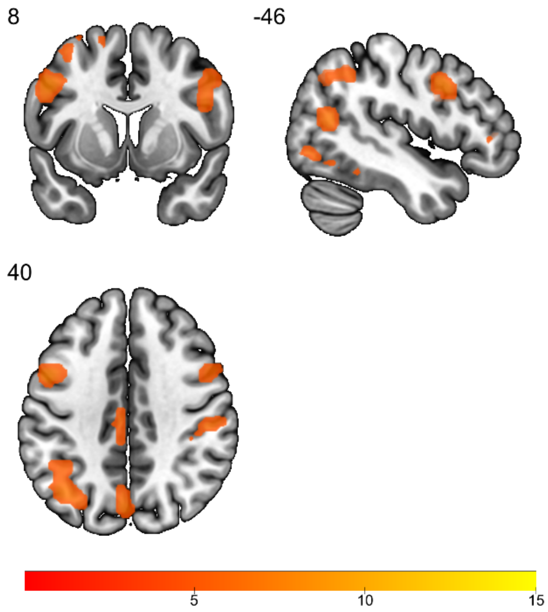


**Figure S6.** DisNeg vs. LookNeg contrast during pre-stimuli anticipation.
